# On the Aging and Reproductive Consequences of Biological Complexity

**DOI:** 10.1101/2022.09.14.505979

**Authors:** Tapash Jay Sarkar

## Abstract

Modeling the dynamics of aging often involves attempting to identify a handful of causal factors and correlating it to some ordered and/or disordered evolution. However, this approach may completely overlook the reality of aging as a distributed and complex evolution. Additionally, the theory driving aging should not just be about the how, but also the why, in that there should be motivated objectives and principles behind growth, decay and alternative routes like reproduction. All of these features are incorporated into a new, theoretical model for the aging process proposed herein. It incorporates a number of techniques from other fields that regularly characterize and predict complex systems dynamics.

## 1. Motivation

Studying the aging process is an effort towards unification. Instead of partitioning and subdividing the individual behaviors of cells, tissue or organs, this field endeavors to understand their common and core dynamics under both healthy and diseased conditions. In establishing this core dynamics, however, the prominent mindset has been very reductionist - trying to isolate the individual or handful of component that fundamentally drives the entire process. For instance, trends like telomeres for explaining cellular population aging or the GH/IGF-1 axis for metabolic aging attempt such simplification. Instead, a more holistic picture looking at and embracing the full complexity of biological systems maybe be much more insightful in detailing how they grow and eventually fail. In fact, this work goes as far as to postulate that increasing system complexity is both the main counter to variety of environmental stressors but also root cause of internal stress from added costs of maintaining that complexity. In addition, biological systems, from cells to species, have an additional unique and characteristic strategy to preserve their complexity, or rather the information encoded within it. This strategy is to reproduce: forgoing the “hardware” manifestation of complexity for the propagation and reinstantiation of the “software” encoding the complexity. This is a critical part of the calculus of biological systems and is weighed in their design and optimization. This directive is unique to biology vs other naturally occurring systems, though some man-made systems try to emulate this, for instance a society and its enduring values or a company and its continuing product line. The story of aging is also intricately interwoven with the story of reproduction, as the former is actually about preserving the hardware while the later is focused on preserving the software and this trade-off is a critical one which will be discussed. Finally and possibly the most fundamental perspective is that biology, like most systems, is not static but instead operates around an equilibrium point. Such points exist because they represent a stable state in which the system can both continue to operate but also be responsive to most internal and external inputs. This optimal steady state is usually reinforced in the design of the system through restorative mechanisms which makes it easy to revert to the state after each perturbation. In medicine this is called the healthy condition, in biology it is called homeostasis. There are of course effects that don’t just perturb but also also evolve the equilibrium point itself, both for greater and for diminished functionality, and it is this evolution of homeostasis which maybe the truest definition of aging itself.

The focus of this paper is to present a simple, yet elegant model that captures these principles and can be empirically utilized to offer insights into the core aspects that characterize the aging and reproductive processes. This model draws from canonical techniques in physics, engineering and optimization theory centered on describing the mechanics of systems with a dynamic equilibrium. This is include the defining an energy or expense function that needs to be minimized. This function also typically incorporates a term that goes as the square of the displacement from equilibrium (the standard spring or restorative potential). The optimal equilibrium state then also typically minimizes the action or accumulated cost over some functional equation, a standard for calculus of variations. In addition, these dynamical systems often involve a recursive mechanics that can often erode the system as a consequence of operation. Additionally, it is a common strategy approximate the mechanics in different operating regimes of the system, for instance a low energy or fast moving regime, or based on time or spatial averaging. These canonical features are now, novelly, all incorporated in the biological systems model proposed herein. Ultimately, it is just a start, but an important one, down the road of a more theoretic and analytic construction of biological evolution with age and reproduction as a dynamical system.

## 2. Model System

The novel model system proposed embodies three core principles. **Principle 1** is the existence of a *Core Complexity Trade-off* between a) increasing the complexity of a biological system to process and counter more diverse environmental stimuli and b) incurring more development costs and failure modes resulting from the increased complexity. **Principle 2** is the occurrence of *Homeostasis-Evolution Defined Life Stages* where programmed growth, stochastic external stimuli and entropic decay all selectively dominate the deviation from the biologic equilibrium. And finally, **Principle 3** is the *Accumulated Cost Motivation for Reproduction* which proposes that nature has optimized the fecundity period of a biological system so that the time-aggregate cost of maintaining the mature, fecund system should not outweigh the cost of producing and maturing a new copy. These Principles most familiarly relate the life of a whole organisms or species but can be applied to the evolution of virtually any biological system – even individual cells, tissues or organs.

Our specific model instantiation of these Principles may be simplistic as it incorporates the widely utilized harmonic oscillator structure around an equilibrium, with additional terms for the magnitude of system complexity and growth. Other, more advanced models maybe more appropriate to model dynamics of specific biological systems, but nevertheless our model captures the core consequences implied by the Principles. It begins with the following equations

The first is the Running Energy/Expense Function

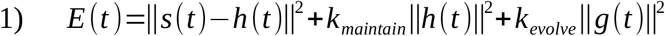

the Evolution of Homeostasis

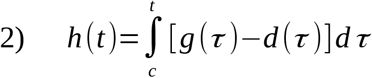

and the Decay Relationship

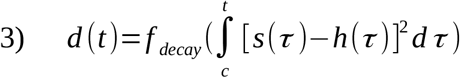

In explaining the model starting with Equation 1, we first note that *E(t)* of course is a the Running Energy/Expense Function that evolves in time *t*. It has a cost for restoring the system after it deviates to state *s* from the biologically equilibrium, otherwise known as the baseline homeostasis *h(t)*. This expense scales with the size of the deviation, as most restorative interactions do. The manifestation of *s(t)* can be any holistic state space but should represent a parameter that is a steady state aspect of the biological system. In addition, each component can be normalized by its own the range of variation so that no component dominates just because it has a greater absolute magnitude. Alternatively, one can also leave them unnormalized to reflect the relative expenses of establishing the larger swing components. An example, a parameter that is constantly cycling, like the proteomic distribution for instance, should be captured by the frequency or rate of production/degradation, which is more steady state, rather than the gross amount of each protein which is constantly fluctuating. The rate can additionally normalized to the average rate of turnover for that protein across multiple instances of the system (e.g. different cells or individuals). In addition there is also a payment to maintaining *h(t)* that scales as some constant *k*_*maintain*_ in this linear penalty model. Finally, there is a payment for growth *g(t)* that scales as some constant *k*_*evolve*_, again in a linear penalty model. Note, one could additionally define a scaling constant *k*_*deviate*_ for the deviation term, however this is just an arbitrary parameter that can be ignored since only the relative scaling between the terms making up *E(t)* matter. Equation 2 describes the Evolution of Homeostasis which integrates the history of the growth *g(t)* as well as a decay term *d(t)*. This captures the idea that the *h(t)* is the culmination of a programmed growth and the erosion of accumulated decay; again arbitrary scaling constants could be added here, but they are instead folded into the definition of *g(t)* and *d(t)*. These terms operate to change the baseline rather than the more realtime evolving *s(t)*, having a more stochastic evolution given external stimuli (environmental factors like food, temperature, physical stimuli etc) and the resulting internal system processing and recovery. Finally, Equation 3 establishes the Decay Relationship for *d(t)* which ties back and incorporates the history of deviations from homeostasis through some decay function *f*_*decay*_. This embodies the principle that the decay of the homeostatic baseline arises from the reality that each excursion from baseline is not perfectly rectified, and instead, leaves some collective, residual impact through *f*_*decay*_. The integral term is the compiling time variance, in a sense, of the state from its equilibrium point, though recall that equilibrium is also evolving with time. The dynamics of *g(t)* could also be detailed, however they are more specific the choice of space and parameter and are thus not focus of this work. Nevertheless *g(t)*’s dynamics can of course be empirically derived, as will be noted a bit later on.

### 2.1 Principle 1: Core Complexity Trade-off

We additionally have the Dimensionality Specifications

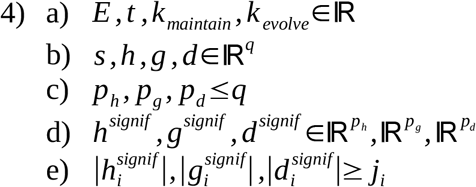

These start to capture the objective of the system which, as with most energy or expense functions, is to reduce *E(t)* over *t* with the weights *k*_*maintain*_ and *k*_*evolve*_, all scalars by Equation 4a. The *q*-dimensional state space is of course defined by the choice in state space. It represents and establishes the overall scope of the parameter being assessed (the total number of proteins, or genes, or behaviors etc) in Equation 4b. However the system may only be designed/trained based on limited subspaces of external and internal stimuli, the dimension sizes *p*_*h*_, *p*_*g*_ and *p*_*d*_ *in* Equation 4c. For instance, some environments involve major swings in temperature – like a dessert – while others alternatively may involve greater variations in the amount of total daylight – like near the poles. The full space of genes may be able to deal with both temperature and total daylight swings, but an organism living in an environment with just the former has no incentive to develop the genes for the later and vice versa. Thus, generally at any point in time the homeostatic baseline may only have significant components in a *p*_*h*_*-*dimensional subspace. At the same time, *g(t)* may be trying to get it to grow into a *p*_*g*_*-*dimensional subspace while *d(t)* may still be storing a history in and trying to erode a *p*_*d*_*-*dimensional subspace. All of this is reflected in Equation 4d. Note, *g(t)* does not just unilaterally increase a component, it can also decrease components as the system evolves and the same versatility is true for *d(t)* as well. The distinction is the *g(t)* is more proactive and thought to make the system better suited for its environment while d*(t)* is reactive and makes it less suitable. Equation 4e explicitly elucidates that significant dimensionality is where the magnitue of the *i*^*th*^ component in the significant subspace is greater than or equal to some arbitrary significance threshold *j*_*i*_, which can of course vary from component to component.

Thus the trade-off optimization proposed by **Principle 1** is manifested in that *g(t)* can work to grow the complexity (significant dimensionality and component size) of *h(t)* to better handle the *q* dimensional stimulus space, however it comes with a price for growth into that space. Yet if *g(t)* is too conservative and *h(t)* and *p*_*h*_ remain too small such that it can’t handle a new stimuli, then not only will it get penalized through *E(t)* but there will be an enduring penalty through *d(t)*. Thus motivating the careful trade-off, fine-tuning and foresight of the biological system based on the likely environments and stimuli it will face. A concrete example could the parameter space of the genome and with the *i*^*th*^ component gene for producing ultraviolet ray resistance protein. A species may have inherited the gene because it originally habituated a sunlight rich environment where it was helpful in establishing a more stable homeostatic baseline. Then perhaps the species migrated to a dense foliage environment with little sunlight. Then the cost of growing and maintaining that UV resistance may not be worth it and the species eventually selects out the gene. However, should there be a drought or a fire or something that clears the sun protection, then the species will be short term penalized through *E(t)*, potentially manifesting as sunburn, and long-term penalized through *d(t)*, potentially manifesting as melanoma, hence the calculated trade-off. Ultimately, the fully mature and optimized baseline *h*^*maturity*^*(t)* should minimize the Deviation Functional *D[h(t)]* in Equation 5 from the conception time *c* to the lifespan *l* over all trajectories of the state space. This is captured by the averaging over all the *s(t)* trajectories that different instances of the system would undergo in that environment.

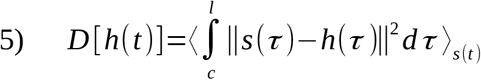

However, without a full knowledge of how the environment will shape the state space for a specific parameter, solving this relationship will be difficult. Instead an empirical derivation of *h*^*maturity*^*(t)* will be proposed later.

### 2.2 Principle 2: Homeostasis-Evolution Defined Life Stages

One can then understand the life stages in **Principle 2** as naturally falling out from different time or age regimes where certain terms dominate.

The first are the Formation Stage Conditions

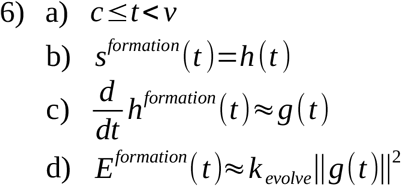

This is the stage from *t* from *c* to viability *v*, denoted in Equation 6a, when the system is not trying to maintain homeostasis but to initially form it and is often supported by a contained environment to establish itself. Thus, there is no sense of deviating from an established baseline due to external stressors, so the system can be seen just at baseline for the duration of the stage, as shown in Equation 6b. This stage has marked growth but no real concept of decay, thus evolution of *h(t)* from Equation 2 reduces to Equation 6c (in derivative form). Then *E(t)* is largely is dominated by the cost of growth in Equation 6d and should generally decrease with time as the system becomes more formed. Thus this stage is driven by pure growth. For an organism as the biological system, this stage would correspond to gestation in the protective and supportive womb or egg environment.

Then the Development Stage Conditions are

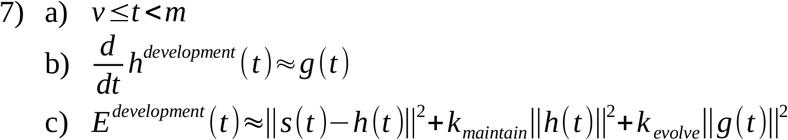

This is the stage from *t* from viability to maturity time *m*, denoted in Equation 7a, when the system is viable, trying to maintain its homeostasis but also trying to grow to meet the demands of the external and internal stimuli. This stage also has substantial growth, though perhaps a lower magnitude compared to the formation stage, but still no real decay, for a similar evolution Equation 7b. Then the *E(t)* has the most expense terms, restoring from environmental challenges, maintaining homeostasis and supporting growth in Equation 7c. The growth will continue to decrease with time, while the state of the system will start to have more fluctuations and costs, but ultimately *E(t)* should continue to decrease as the system becomes more optimized and finally matures. Thus this stage reflects a mix of growth and interaction with the environment. For the organism example, this would be like childhood and adolescence.

Following that, are the Maturity Stage Conditions

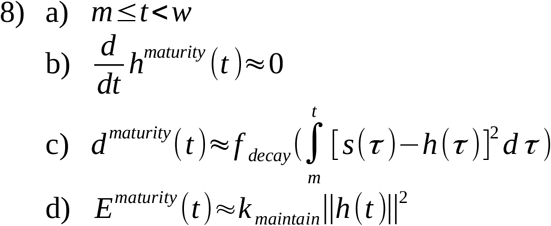

This is the stage from *t* from maturation to the waning point *w*, denoted in Equation 8a, when the system has achieved its optimal state of reduced running expenses. The growth has now ceased as the system has reached its optimum and the decay term is small enough so that *h*^*maturity*^ largely remains the same, in Equation 8b. That being said, the decay term is starting to accumulate now through the *f*_*decay*_ function in Equation 8c and will be relevant for the next stage. However, for this stage, since *h(t)* is relatively constant and *h*^*maturity*^ minimizes the deviation over all the trajectories *s(t)* (established in Equation 5), the deviation term can be assumed to be negligible for *E(t)* with just the maintenance cost dominating Equation 8c. Thus this stage represents an inertial phase, where the cost is minimized and there is neither growth nor decay, allowing the system to coast in its optimal state for a duration. For the organism example, this would be adulthood.

Finally, there are the Senescence Stage Conditions

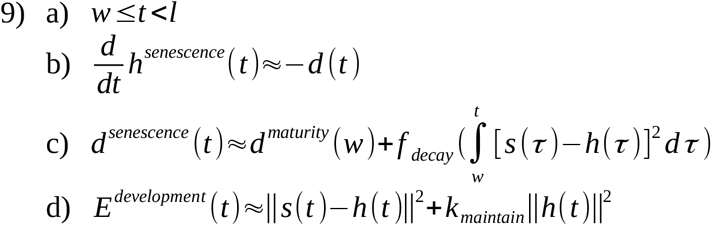

This is the stage from *t* from waning to the end of lifespan *l*, denoted in Equation 9a, when the system is now realizing the accumulated operation and deviation expenses while still trying to cope with new environmental stimuli. In this stage, the decay has now grown significant that it can no longer be ignored, in Equation 9b and Equation 9c. It moves the system out of the inertial phase and once again causes *h(t)* to deviate from baseline more substantially and significantly contribute to *E(t)* in Equation 9d. This phase is like the frictional or entropic erosion slowing the inertial free motion and taking over the system, now that there is no driving growth element. For the organism example, this would be be middle and old age, with the waning point also potentially corresponding to a menopause or andropause type point.

Additionally we have the ultimate Viability Threshold

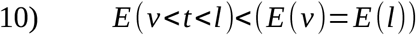

Equation 10 holds that the cost of the viable system must always be less than that at the viability point or equivalently the end of lifespan point. Of course before viability the system is forming and after lifespan it is dead, so this is just threshold of life.

### 2.3 Principle 3: Accumulated Cost Motivation for Reproduction

Equation 11 establishes the concept of Total Action/Accumulated Cost incurred by the system as defined as

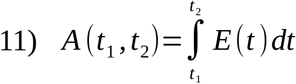

This is distinct from the decay term, which only factors in the expenses of deviating from homeostasis. This then sets the framework for the Reproduction Justification in **Principle 3** in that, once it is more expensive to continue maintaining the mature system than to form and develop a new one, then it is more optimal to reproduce. Thus the system is designed such that the maturity stage accumulated cost that should not outweigh the formation and development stages accumulated cost (of course in general, since each system will take its own unique trajectory). So the two should be equal when averaging over all state space trajectories l as captured in Equation 12

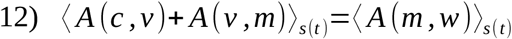

This relationship can be used to back out the average length of the development stage and the maturity stage, given that the formation duration for a biological system is typically already known. The senescence phase is not directly bounded by *A(w,l)*, but is terminated instead upon *E(l)* crossing the viability threshold. However since it has a dominant decay term, it has it’s own implicit accumulation of costs which is like an indirect, deviation specific, accumulated cost constraint. The average lifespan is typically known, though could be calculated from these dynamics.

These relations establish the model and meet the **Principles**. In the next section, the empirical reduction and application of the model will be discussed.

## 3. Empirical Reductions

In the previous section, a number of stage specific reductions were described, which help simplify the calculations. However for the empirical application of modeling the system behavior, a number of additional simplifications and implementation choices can be made.

First, it can be noted that *s(t)* can of course just be empirically measured as the state of the system at different time points. However the homeostatic baseline is more of a hidden variable. However an Empirical Baseline can be established as

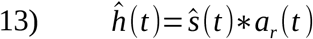

where *a*_*r*_*(t)* in Equation 13 represents some window averaging function of *ŝ(t)* to estimate *ĥ (t)*. This function operates over a range *r* in time, such that the convolution with *a*_*r*_ conveys the construct of age as linked to the average (homeostatic) state when binning or smoothing over windows of *r* units of time. Then by the properties of convolution and the stage specific reductions we have the Empirical Dynamics

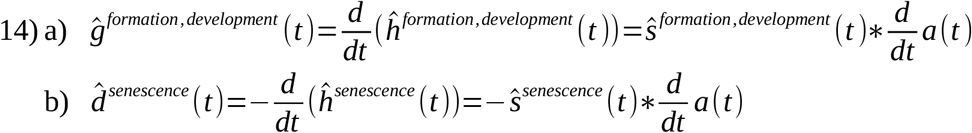

where we can calculate the dynamics based on the change in the smoothed state function or convolve it with the derivative of the window averaging function in Equation 14a and Equation14b.

A simple starter averaging function would be the rectangle function of width *r* and height 1*/r* so it is more of a square function, or *П*_*r*_. Though, for the sake of differentiability, instead *á*_*r*_*(t)* can be defined as

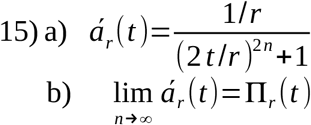

since the limit of Equation 15a is the *П*_*r*_ square function in Equation 15b. Then the derivative function becomes

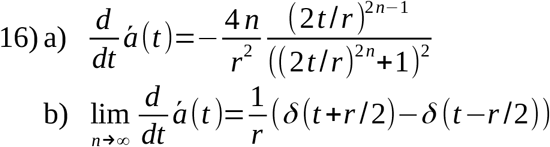

since the limit of Equation 16a is the scaled difference of two delta function in square function in Equation 16b. This then reduces the variable set to Boxcar Averaged Set

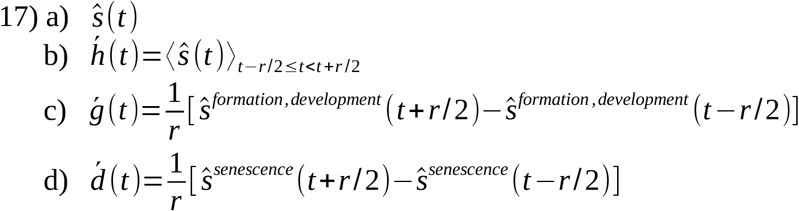

which shows that all the variables in the model can be calculated from the time series of *ŝ(t)* in Equation 17a from Equation 17b, Equation 17c and Equation 17d. Furthermore we can considered a modified Expense Function that also smooths the cost by the same function *á*_*r*_*(t)*

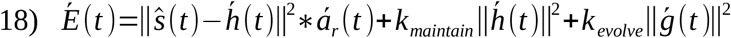

where the only smoothed term in Equation 18 is the deviation contribution, as the other terms are already based on smoothed variables.

This gives us a smoother payment function *É*to satisfy the Reproduction Justification Equation 12 with the general behavior depicted in Figure 1.

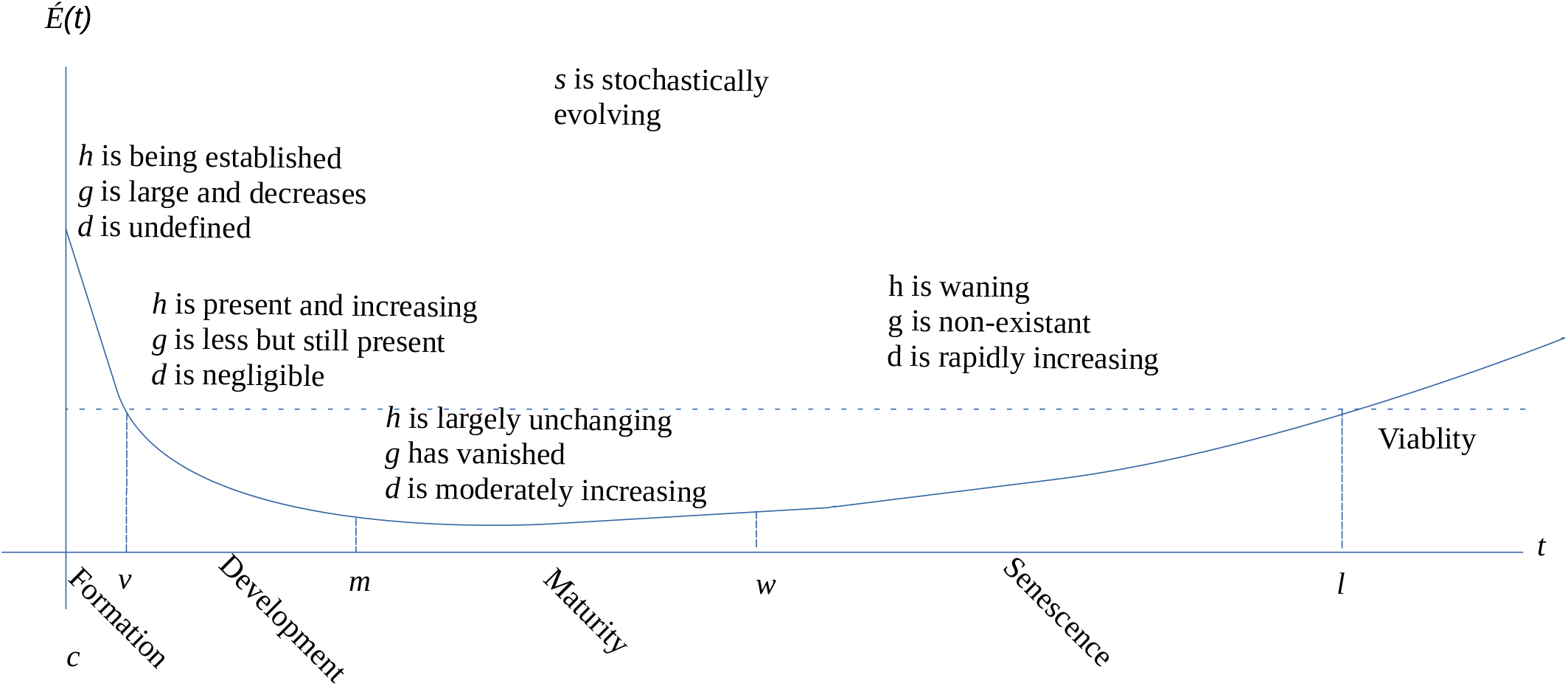

## 4. Directions

With the reductions in place there are a number of consequences that can be explored with the model for any specific biological system it is applied to.

First of course is the significant window *r* with which the dynamics scale. Any state space time course can be smoothed with multiple such scales and the time scale that best fits the data to all Stage Dependent Conditions and the Reproduction Justification could unearth something about the true aging rate of the system. Even more intriguing, a more dynamic smoothing function could also be used, for instance one that changes in each stage or even that is continually, gradually changing. This could reflect the fact that the system may age at different rates throughout its lifespan and better characterize and quantify biological age and rate vs chronological age. This can maybe even be made personalized to each instance of the system, as long as there is sufficient time course data; this would truly demonstrate that the influences of each state space trajectory as influenced by the specific environment history is reflected in the system’s core evolution and specific aging process.

Second, as alluded to before, the duration of the various life stages of the biological system can be determined from the evolution of the homeostatic function. Though these are estimated empirically, a more natural delineation of the phases, in particular of the *m* and *w* time points, may fall out of the time course data when applied with the model constraints. The onset of maturity is currently defined by single features, for instance the development of a reproductive component or the optimization of a particular function like mechanical force output. However a more comprehensive picture of maturity involving the full complexity of some parameter space may be more telling of objectives of the system. Then understanding the reproductive or replication window in relation to the full complexity of maturity can better define when the system truly starts waning. This could be used to instruct optimal points and aspects for intervention. In addition it could explain an evolutionary picture behind the system if differentially evolved instantiations of the systems exist and can be compared, like the varying genomic state space complexity between different species in a phylogentic tree. The differing windows of each life stage can be correlated with the complexity of the species and the complexity of the habitat it survives in. Perhaps, even species that cohabitate an environment or the same species in different environments can be compared under the model fitting to suss out the independent contributions of the two sources of complexity (nature vs nurture), in sort of partial derivative or controlled metadata analysis.

Third is the empirical mapping out of *f*_*decay*_. This function details the dynamics of the advanced aging process as it interprets and registers deviations from homeostasis as accumulated stressors. Trying to understand the dynamics and especially the causality behind the advanced aging process in organisms is the core focus of the field of longevity research. The shape of this function, established through fitting the model, and possibly even its own evolution with time, would be very impactful for this research. Furthermore the multi-dimensionality of this function could bring the field closer to a true understanding of the advanced aging process of an organism or other biological systems in general. As mentioned before, the reductive approach often taken when characterizing these dynamics is often not only lacking comprehensiveness or correlation, but it also may completely miss the causality of the process. Specifically, they miss the rationale that the advanced aging rate and mechanics may be in line with and could be seen as consequence of achieving more complex and robust systems to handle an environment. Thus tying an evolutionary imperative with the designed system longevity.

Fourth is the question of what are the most natural parameter spaces that embody these dynamics. The ones that are comprehensive enough and best fit the model construct may be key parameters that drive the growth and lifetime of the system. A natural place to look would be parameters that have already been identified for algorithmic clocks, for instance the case of DNA methylomics. The methylome is the sequence of chemically modified cytosine nucleobases in the cellular genome. They defines the first layer of what is known as the epigenome – the architecture that does not change a gene’s coding but instead its activity. High frequencies of CG sequences, known as CpG sites, tend to populate the promoter regions of a gene. These are the regions that transcription machinery searches for and binds to to initiate transcription of that gene into mRNA. Thus, the addition of the methyl group H_3_C to the pyramadine rings of cyotsine at a CpG site can prevent the binding of this transcription machinery. In addition, this methylation can engage the binding of methyl-CpG-binding domain proteins which further repress gene expression through more complex epigenetic features like histone modifications and chromatin compaction. Thus cytosine methylation often correlates with gene silencing. Much work has been recently done building linear correlative models connecting tissue age with the bulk methylation pattern in its cells, with relatively high accuracy in a variety of tissues. These models specifically use a parameters called *β*_*i*_ values, which is the percent of all copies of the i^th^ CpG site that are methylated over the cells analyzed in a tissue. This would be an easy and natural starting application for the model posed here as methylation, epigenetics in general, are a more steady state space which still evolves over time. Furthermore, the *β* values are normalized to the total number of copies of each site, so choosing them as the state vector *s(t)* would be an equal weighting of all the sites in the cost evaluations. This presents an interesting implementation, though an unequal weighting could factor in the additive cost each methylation event. And of course there are multiple “omics” parameters spaces that could also be good fits, and perhaps the best fit would constitute the most natural lens by which to view the aging process.

Ultimately this model is unique in that it is the first to introduce a number of imperatives in its design, with its Principles on the Core Complexity Trade-off, Homeostasis-Evolution Life Stages, Reproductive Justification and its nature as full Multi-Variate Model. It is of course a simple, mostly linear model but it opens up the discussion and investigation into the tremendous value of holistic and first principles based understanding to the aging process.

